# Characterization of neutralizing antibodies from a SARS-CoV-2 infected individual

**DOI:** 10.1101/2020.05.12.091298

**Authors:** Emilie Seydoux, Leah J. Homad, Anna J. MacCamy, K. Rachael Parks, Nicholas K. Hurlburt, Madeleine F. Jennewein, Nicholas R. Akins, Andrew B. Stuart, Yu-Hsin Wan, Junli Feng, Rachael E. Nelson, Suruchi Singh, Kristen W. Cohen, M. Juliana McElrath, Janet A. Englund, Helen Y. Chu, Marie Pancera, Andrew T. McGuire, Leonidas Stamatatos

## Abstract

B cells specific for the SARS-CoV-2 S envelope glycoprotein spike were isolated from a COVID-19-infected subject using a stabilized spike-derived ectodomain (S2P) twenty-one days post-infection. Forty-four S2P-specific monoclonal antibodies were generated, three of which bound to the receptor binding domain (RBD). The antibodies were minimally mutated from germline and were derived from different B cell lineages. Only two antibodies displayed neutralizing activity against SARS-CoV-2 pseudo-virus. The most potent antibody bound the RBD in a manner that prevented binding to the ACE2 receptor, while the other bound outside the RBD. Our study indicates that the majority of antibodies against the viral envelope spike that were generated during the first weeks of COVID-19 infection are non-neutralizing and target epitopes outside the RBD. Antibodies that disrupt the SARS-CoV-2 spike-ACE2 interaction can potently neutralize the virus without undergoing extensive maturation. Such antibodies have potential preventive/therapeutic potential and can serve as templates for vaccine-design.

**IN BRIEF:** SARS-CoV-2 infection leads to expansion of diverse B cells clones against the viral spike glycoprotein (S). The antibodies bind S with high affinity despite being minimally mutated. Thus, the development of neutralizing antibody responses by vaccination will require the activation of certain naïve B cells without requiring extensive somatic mutation.

**Highlights:**

- Analysis of early B cell response to SARS-CoV-2 spike protein
- Most antibodies target non-neutralizing epitopes
- Potent neutralizing mAb blocks the interaction of the S protein with ACE2
- Neutralizing antibodies are minimally mutated

## INTRODUCTION

The WHO declared the 2020 COVID-19 to be a global pandemic on March 11^th^, 2020 (World Health Organization, 2020). There are currently 4.2 million documented cases of COVID-19 and over 290 000 deaths (Dong et al., 2020). The infection is caused by SARS-CoV-2, a beta coronavirus, closely related to SARS-CoV (Wan et al., 2020). Presently the immune response to COVID-19 is not well understood and preventative measures, such as vaccines, are not available. It is also unclear which immune responses are required to prevent or control SARS CoV-2 infection.

High resolution structures of the SARS-CoV-2 prefusion-stabilized spike (S) ectodomain revealed that it adopts multiple conformations with either one receptor binding domain (RBD) in the “up” or “open” conformation or all RBDs in the “down” or “closed” conformation, similar to previous reports on both SARS-CoV S and MERS-CoV S (Gui et al., 2017; Kirchdoerfer et al., 2018; Pallesen et al., 2017; Song et al., 2018; Walls et al., 2020; Walls et al., 2019; Wrapp et al., 2020; Yuan et al., 2017). Like SARS-CoV, SARS-CoV-2 utilizes angiotensin-converting enzyme 2 (ACE2) as an entry receptor binding with nM affinity (Li et al., 2003; Walls et al., 2020; Wrapp et al., 2020) (Hoffmann et al., 2020; Letko et al., 2020; Ou et al., 2020). Indeed, the S proteins of the two viruses share a high degree of amino acid sequence homology; 76% overall and 74% in RBD (Wan et al., 2020).

Although binding and neutralizing antibody responses are known to develop following SARS-CoV-2 infection (Ni et al.; Okba et al., 2020), no information is currently available on the epitope specificities, clonality, binding affinities and neutralizing potentials of the antibody response.

Monoclonal antibodies (mAbs) isolated from SARS-CoV-infected subjects can recognize the SARS-CoV-2 S protein (Yuan et al., 2020) and immunization with SARS S protein can elicit anti-SARS-CoV-2 neutralizing antibodies in wildtype, and humanized mice, as well as llamas (Walls et al., 2020; Wang et al., 2020; Wrapp et al., 2020). However, SARS-CoV-2 infection appears to not elicit strong anti-SARS-CoV neutralizing antibody responses and vice versa (Ou et al., 2020).

Here, we employed diverse but complementary approaches to investigate the serum binding and neutralizing antibody responses to a stabilized ectodomain variant of the SARS-CoV-2 spike protein (S2P)as well as the frequency and clonality of S2P-specifc B cells in a SARS-CoV-2-infected individual 21 days post post the onset of respiratory symptoms. We isolated anti-SARS-CoV-2 S mAbs and characterized their binding properties and determined their neutralizing potencies. Among all B cells analyzed, no particular VH or VL gene family was expanded and the isolated antibodies were minimally mutated. Our analysis reveals that only a small fraction of S2P-specific B cells recognized the RBD. Of the forty-four mAbs analyzed, only two displayed neutralizing activity. The most potent mAb, CV30, bound the RBD in a manner that disrupted the spike-ACE2 interaction. The second mAb, CV1, bound to an epitope distinct from the RBD and was much less potent.

## RESULTS

### Serology

Serum and PBMC were collected twenty-one days after the onset of clinical disease. The serum contained high titers of antibodies to the SARS-CoV-2 S2P (Fig. 1A). The specificity of this response was confirmed by the absence of S2P reactivity by serum antibodies isolated from donors collected prior to the SARS-CoV-2 pandemic, or donors with confirmed infection by endemic coronaviruses. We also measured the serum antibody response to RBD, and again observed specific high titers of binding antibodies (Fig. 1B). Isotype-specific ELISA revealed that the IgG titers were higher than the IgA and the IgM titers to both S2P and RBD, suggesting a significant portion of the antibody responses to SARS-CoV-2 S are IgG (Fig. 1C and D). The serum from the SARS-CoV-2 infected donor displayed potent neutralizing activity (Reciprocal ID_50_∼3000)against a pseudovirus expressing the S protein from SARS-CoV-2 isolate Wuhan-Hu-1 (Fig 1E). We concluded that this donor had developed strong binding and neutralizing antibody responses within three weeks of disease onset.

**Figure 1.**
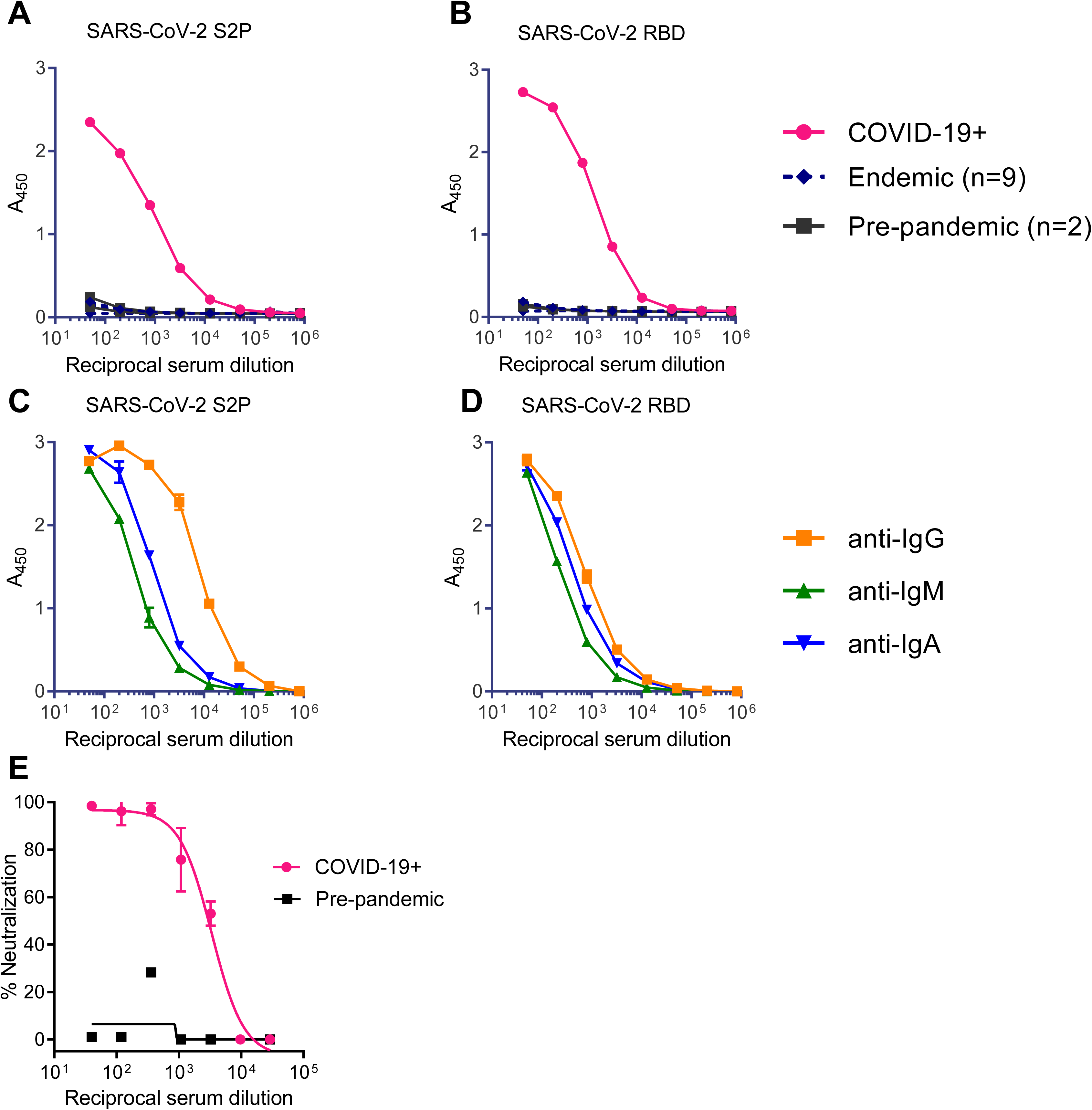
Serum antibody reactivity to the SARS-CoV-2 ecto- and receptor binding domain. Total antibody binding in serum from a donor with confirmed SARS-CoV-2 infection (COVID-19+), from two donors collected prior to the COVID-19 pandemic with an unknown history of coronavirus infection (pre-pandemic), and from nine donors with confirmed infection by endemic corona viruses (endemic), was tested for binding to the SARS-CoV-2 S2P ectodomain (**A**) and the RBD (**B**) by ELISA. Serum from the donor in SARS-CoV-2 infection in **A** was tested for binding to the SARS-CoV-2 S2P ectodomain (**C**) and the RBD (**D**) using isotype-specific secondary antibodies by ELISA. (**E**) Serum from donor with confirmed SARS-CoV-2 infection, and serum from a pre-pandemic donor were evaluated for their ability to neutralize a SARS-CoV-2 pseudovirus.

### B cell sorts and VH/VL sequencing

Fluorescently labeled S2P and RBD probes were used as baits to identify B cells specific to the SARS-CoV-2 S protein that were circulating at this timepoint. S2P was labeled with either phycoerythrin (PE) or brilliant violet 711 (BV711) and used to stain B cells concurrently. This double labeling strategy helps to discriminate between bona fide S2P-specific B cells and non-specific background staining to the fluorophores. RBD was labeled with alexa fluor 647 to identify B cells specific for that domain.

Approximately 0.65% of total CD19+ B cells were S2P positive compared to 0.07% of total B cells from a naïve donor (Fig. S1A and B). The dominant responding B cells were IgM+ IgD+ (49% of S2P+ B cells; Fig. S1C); 90% of which were CD27+ suggesting that although these B cells have not class-switched, they were antigen-experienced memory B cells. The second most prominent subset of S2P-specific B cells were class-switched IgG+ IgD-B cells (27% of S2P+ B cells; Fig. S1C). In fact, 1.7% of the IgG+ B cells stained with S2P (Fig. 2A) and of those ∼7% (or 0.12% of total IgG+ B cells, Fig. 2B) were also positive for RBD.

**Figure 2.**
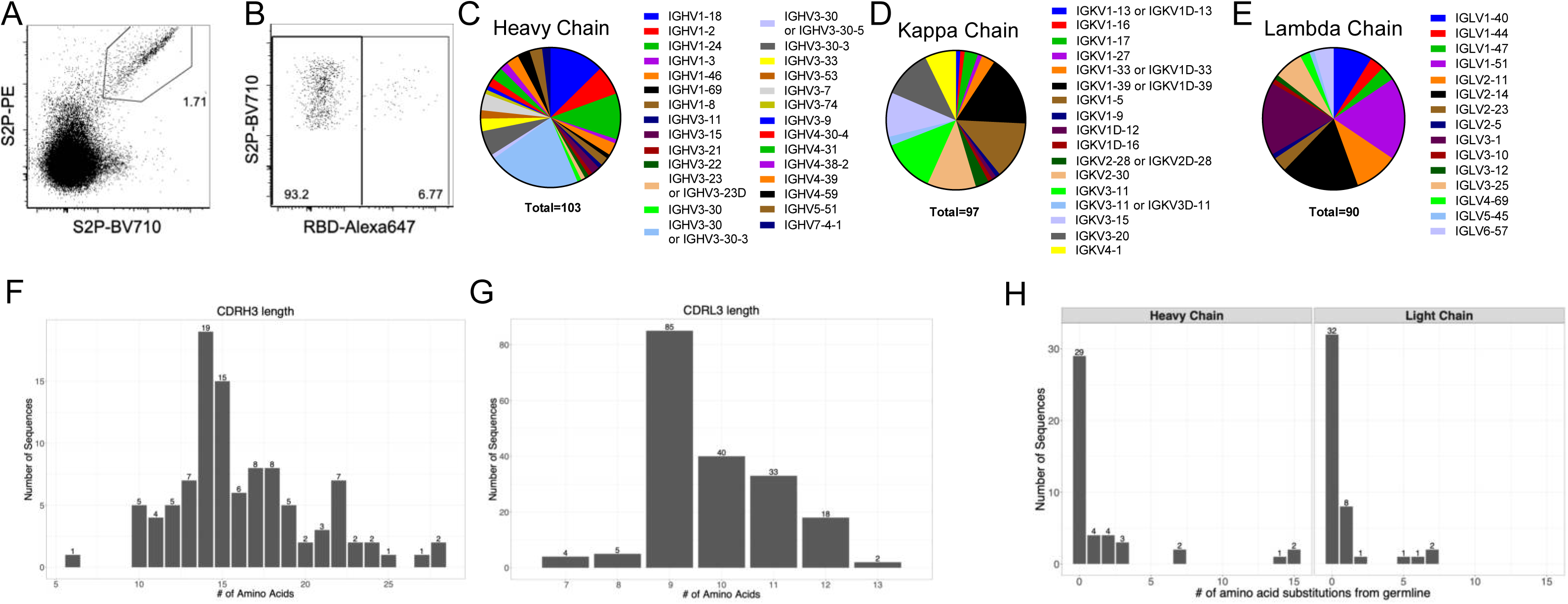
Early B cells response to SARS-CoV-2 is diverse and largely unmutated. (**A**) Class switched (IgM-IgG+) B cells were stained with SARS-CoV-2 S2P labeled with BV710 or PE. **(B)** SARS-CoV-2 S2P+ IgG+ B cells were further analyzed for binding to Alexafluor647-labeled SARS-CoV-2 RBD. (**C, D, E**) Individual SARS-CoV-2 S2P+ IgG+ B cells were sorted into separate wells of a 96 well plate and sequenced using RT-PCR. VH (**C**), VK (**D**), and VL (**E**) gene usage of successfully sequenced S2P-specific B cells. CDRH3 (**F**) and CDRL3 (**G**) length distributions of successfully sequenced S2P-specific B cells. Number of amino acid substitutions from germline in S2P-specific heavy and light chains (**H**).

We hypothesized that the class-switched SARS-CoV-2-specific B cells were more likely to have undergone some affinity maturation and contain antibodies capable of neutralizing the virus. Thus, we focused on our B cell receptor (BCR) sequencing on S2P+ IgG+ B cells. 576 S2P+ B cells were single-cell sorted into individual wells of a 96 well plate and the variable heavy and light chain regions of B cell receptor transcripts were sequenced using nested RT-PCR. We successfully recovered 103 successful VH sequences, and 187 successful VL sequences, 97 of which were kappa and 90 were lambda. B cells specific for S2P+ were derived from diverse antibody heavy and light chain genes (Fig. 2C-E) and had normal distributions of CDRH3 and CDRL3 lengths (Fig. 2 F-G). Consistent with the relatively short time of infection the majority of BCR sequences showed low levels of somatic mutation (Fig. 2H).

### Antibody-binding

Among all successfully sequenced VH and VL transcripts, we obtained paired sequences from forty-four. These were produced as recombinant monoclonal antibodies (mAbs) of the IgG1 isotype and tested for binding to recombinant S-derived proteins (Fig. 3).

**Figure 3.**
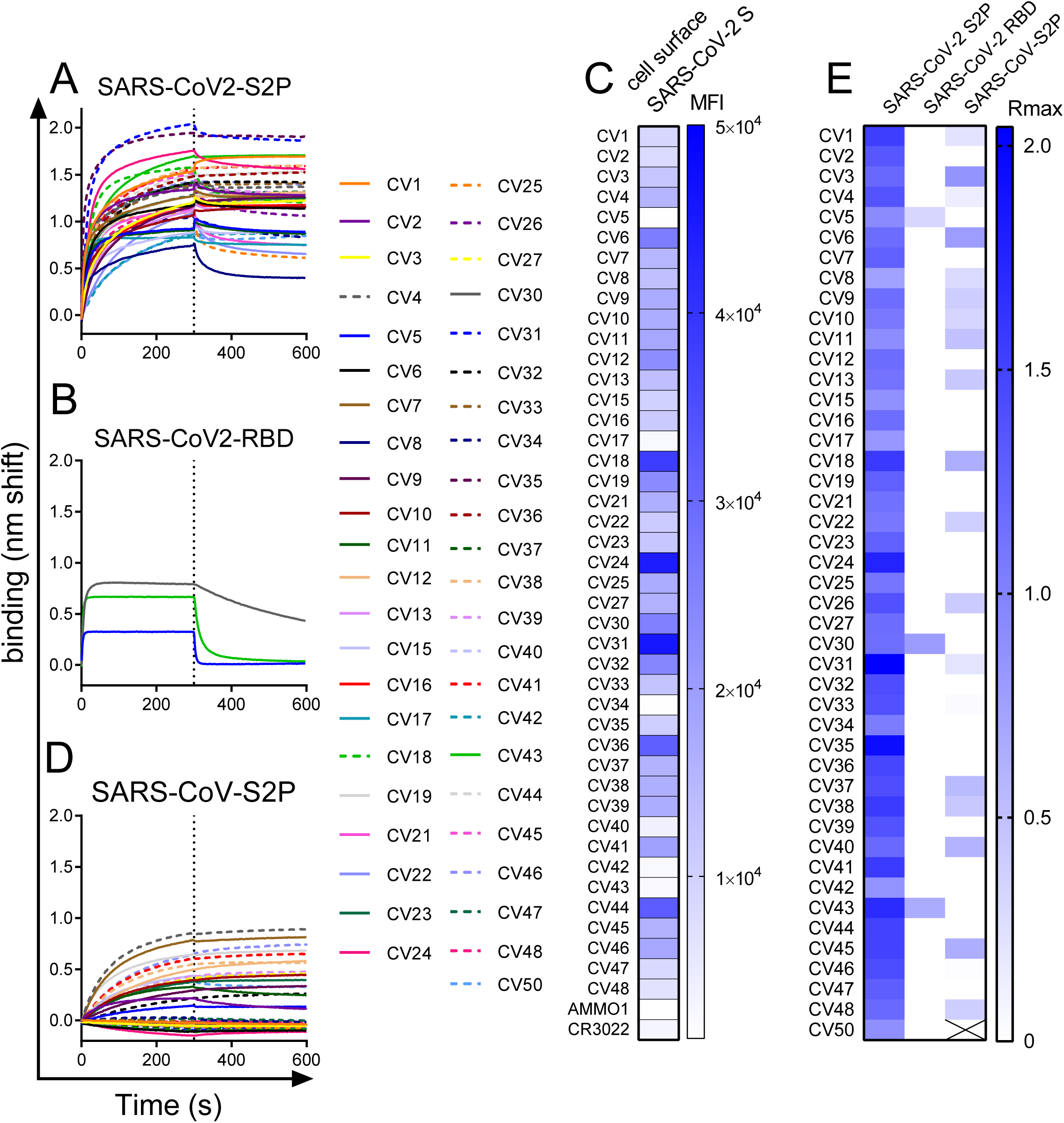
Sorted mAbs bind to SARS-CoV-2 and a subset cross-react with SARS-CoV S. mAbs isolated from SARS-CoV-2 S2P-specific B cells were tested for binding to SARS-CoV-2 S2P (**A**) and to SARS-CoV-2 RBD (**B**) using BLI. (**C**) mAbs were labeled with phycoerythrin (PE) and used to stain 293 cells transfected with wildtype SARS-Cov-2 S by flow cytometry. Heatmap shows mean fluorescence intensity of PE+ cells at 2.5µg/ml. Titration curves are shown in Fig. S2. (**D**) mAbs were tested for binding to SARS-CoV S2P by BLI (**D**). (**E**) Heatmap shows maximum binding response (average nm shift of the last 5 seconds of association phase) of binding data in **A**, **B** and **D**.

All the mAbs bound to the stabilized SARS-CoV-2 ectodomain, S2P (Fig. 3A and E). Consistent with the B cell staining that revealed very few RBD-specific B cells (Fig. 2B), only three mAbs, CV5, CV30 and CV43, also bound the SARS-CoV-2 RBD (Fig. 3B and E). The majority of S2P-specific mAbs also bound to full-length membrane-bound wildtype SARS-CoV-2 S on the surface of 293 cells (Fig. 3C and Fig. S2). The observation that some S2P-specific mAbs failed to bind to cell surface S indicates that there may be conformational differences between the stabilized soluble ectodomain and cell surface S. The fact that a subset of the mAbs bound to a stabilized ectodomain variant of the closely related SARS-CoV S protein (Fig. 3D and E), demonstrates that there are conserved epitopes among the two viruses. Consistent with the lower degree of conservation of the S1 subunit between SARS-CoV and SARS-CoV-2, the anti-RBD mAbs CV30 and CV43 did not cross react with SARS-CoV S2P, while CV5 showed weak binding.

### Neutralizing activity

The S2P-binding mAbs were evaluated for their ability to neutralize SARS CoV-2 pseudovirus infection of 293T cells stably expressing ACE2. All but two of the mAbs were non-neutralizing (Fig. 4A and Table S1). Although it did not achieve 100% neutralization at the highest concentration, CV1 which binds an epitope outside the RBD was weakly neutralizing (IC_50_=15µg/ml, Fig. 4A and Table S1). CV1 neutralized less potently than an ACE2-Fc fusion protein which acts as a soluble competitor for the interaction between S and the cell surface-expressed ACE2 (IC_50_=2.2 µg/ml). In contrast, CV30 achieved 100% neutralization and was ∼480 times more potent than CV1 (IC_50_=0.03µg/ml, Fig. 4A and Table S1). CV1, CV30 and the ACE2-Fc fusion did not neutralize a murine leukemia virus pseudovirus demonstrating their specificity for the SARS-CoV-2 S protein (Fig. 4B). CV30 is derived from a heavy chain utilizing an IGHV3-53*01 heavy chain and an IGKV3-30*01 light chain. CV1 binds an epitope outside the RBD and is derived from an IGHV4-38*02 heavy chain and an IGLV1-44*01 light chain. Both represent unique clones among all B cells sequenced (Table S1).

**Figure 4.**
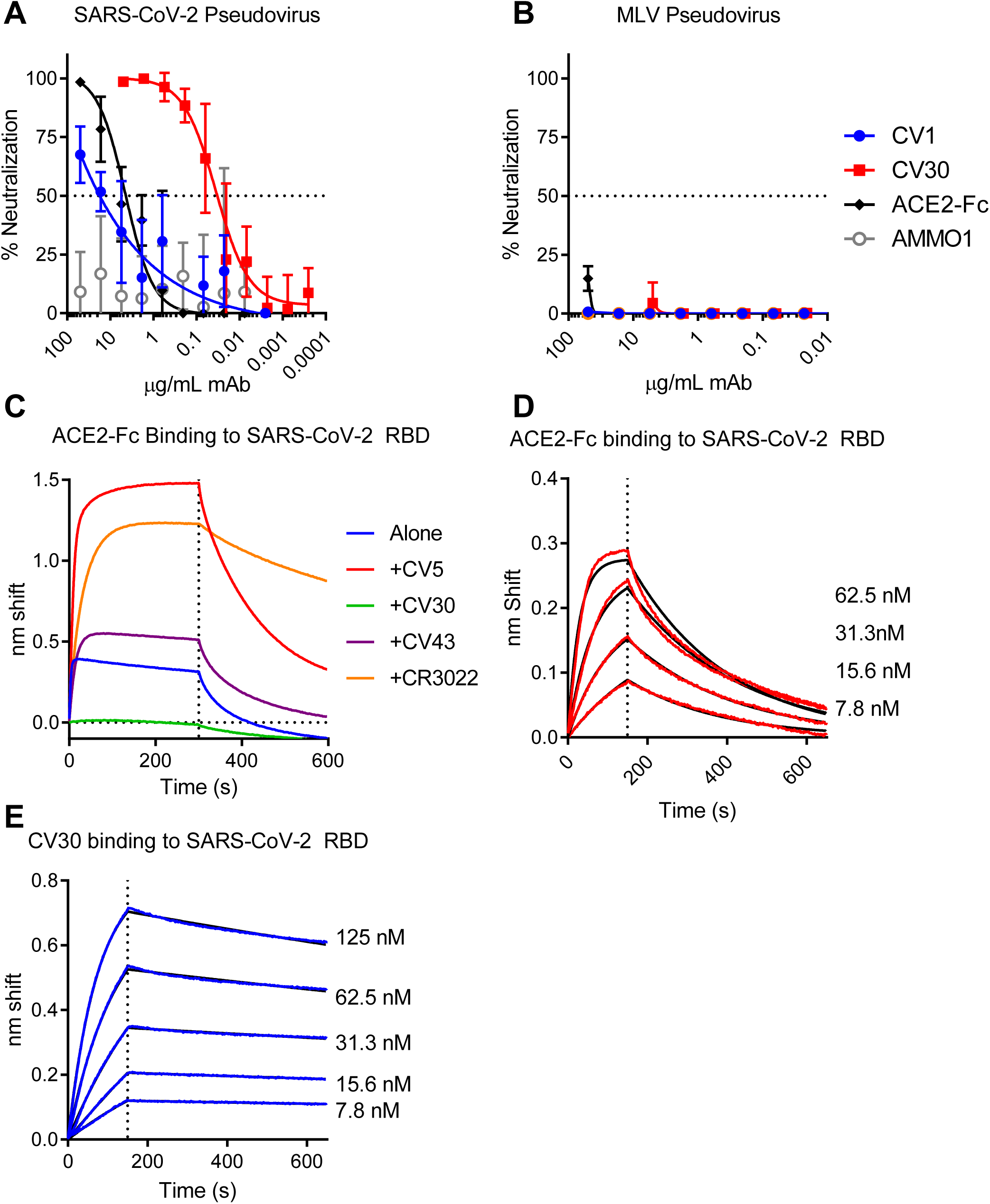
The RBD-specific mAb CV30 neutralizes SARS CoV-2 by blocking the ACE2-SARS-CoV-2 S interaction. (**A**) CV1 and CV30 were serially diluted and tested for their ability to neutralize SARS-CoV-2 pseudovirus infection of 293T cells stably expressing ACE2. An ACE2-FC fusion and the anti-EBV mAb AMMO1 were included as positive and negative controls. Data are representative of 6 independent experiments (see Table S1 for details). (**B**) The same mAbs were tested for neutralization of an MLV pseudovirus. (**C**) Biotinlyated ACE2-Fc was immobilized on streptavidin biosensors and then tested for binding to SARS-CoV-2 RBD in the absence and presence of the indicated mAbs using BLI. (**D**) ACE2-Fc was immobilized Protein A biosensors and binding to the indicated serial dilutions of SARS-CoV-2 RBD were measured by BLI and used to determine the binding constant (kD). Red lines represent the measured data and black lines indicate the theoretical fit. (**E**) CV30 was immobilized onto anti-human Fc biosensors and binding to the indicated serial dilutions of SARS-CoV-2 RBD were measured by BLI and used to determine the binding constant (kD). Blue lines represent the measured data and black lines indicate the theoretical fit. Kinetic measurements from **D** and **E** are summarized in Table S2.

Based on the observations that CV30 is potently neutralizing and it binds RBD with high affinity, we investigated whether it would block the interaction between the SARS-CoV-2 protein and the ACE2 receptor. To this end we setup binding competition experiments using BLI. Indeed, CV30 completely inhibited the RBD-ACE2 interaction. In contrast, CV5 and CV43, the other two anti-RBD mAbs and the CR3022 control, which binds the RBD outside of the binding site, did not (Fig. 4C). We also measured the relative binding affinities of CV30 and ACE2 to the SARS-CoV-2 RBD. ACE2 bound the RBD with an affinity of 5.9 nM (Fig. 4D), while CV30 bound with a slightly higher affinity of 3.6 nM (Fig. 4E). The kinetics of the interactions were notably different, ACE2 had both a faster association and dissociation rate than CV30 (Figs. 4D and E, and Table S2). Collectively these results indicate that CV30 neutralizes SARS-CoV-2 infection by blocking the S-ACE2 interaction through an interaction that is higher affinity.

## DISCUSSION

The development of therapeutic interventions, of immunoprophylaxis and of an effective vaccine against SARS-CoV-2 will benefit from understanding the protective immune responses elicited during infection. A recent report indicated that neutralizing antibodies are present in the sera collected from convalescent COVID-19 patients (Ni et al.; Okba et al., 2020). However, the kinetics of neutralizing antibody development as well as the characteristics and epitope specificities of neutralizing antibodies generated during SARS-CoV-2 infection are presently poorly understood.

Serological analysis revealed that this COVID-19-infected patient developed high titers of binding and neutralizing antibody responses twenty-one days following infection. The development of neutralizing antibody titers at this early timepoint has been reported for other COVID patients (Ni et al.; Okba et al., 2020) and is consistent with the rapid development of neutralizing responses to SARS-CoV infection (Corti et al., 2011). At this time point, IgG constituted the major fraction of anti-S2P and anti-RBD serum antibodies, although both IgM and IgA antibodies against these viral antigens were detected in the serum as well. The S2P-specific, class switched B cells circulating at this time point were not dominated by any particular clone. Rather, they were derived from a diverse VH/VL gene repertoire, with frequencies similar to those reported in healthy uninfected individuals (Briney et al., 2019; DeKosky et al., 2016; Soto et al., 2019; Vazquez Bernat et al., 2019).

Although anti-S2P antibodies isolated at this time point could bind the S protein, the majority lacked neutralizing activity. However, two, CV1 and CV30, were able to neutralize SARS-CoV-2. CV1 and C30 were derived from unique rearrangements among all antibodies we examined. Thus, although diverse B cell clones became activated during infection, the serum neutralizing activity is due to a relatively small subset.

CV1 binds to an unknown epitope region outside of the RBD, but the more potent CV30 recognizes the RBD and likely neutralizes infection by directly inhibiting SARS-CoV-2 S binding to the ACE2 receptor. The RBD is a major target of neutralizing antibodies in SARS-CoV infection (Cao et al., 2010). Several neutralizing monoclonal antibodies that block the interaction of SARS-CoV with the ACE2 receptor have been described (Hwang et al., 2006; Prabakaran et al., 2006; Rockx et al., 2008; Sui et al., 2004; Walls et al., 2019). Moreover, the neutralizing potency correlated with the degree of S-ACE2 inhibition (Rockx et al., 2008). The RBD of MERS-CoV is also a target of potent neutralizing antibodies (Jiang et al., 2014; Niu et al., 2018; Tang et al., 2014; Ying et al., 2014), highlighting the importance of receptor blocking antibodies for coronavirus vaccine development. Although SARS-CoV and SARS-CoV-2 share extensive amino acid sequence in the receptor binding domain (74%) and both viruses utilize human ACE2 for entry, the amino acid identity in the receptor binding motif is only ∼50% (Wan et al., 2020). In line with this, potent anti-SARS-CoV neutralizing monoclonal antibodies that bind RBD fail to cross react with SARS-CoV-2 (Wrapp et al., 2020), similarly the anti-RBD mAb, CV30 described herein fails to cross react with the SARS-CoV spike protein.

Consistent with the short time period post-infection, the majority of S-specific BCRs from individual B cells were unmutated or had only accumulated very few mutations. This was true for the neutralizing antibodies as well. CV1 was unmutated from germline, while CV30 had 2 amino acid mutations in VH and none in VL. Largely unmutated antibodies against SARS-CoV S (Prabakaran et al., 2006; Sui et al., 2004) and MERS-CoV S (Jiang et al., 2014; Tang et al., 2014; Ying et al., 2014; Ying et al., 2015) have been isolated from phage display libraries created from uninfected donors.

Potent anti-SARS-CoV neutralizing monoclonal antibodies that bind RBD are derived from different VH genes (VH1-18, VH1-69, or VH3-30) than CV30 (Prabakaran et al., 2006; Sui et al., 2004; Traggiai et al., 2004; Walls et al., 2019). Anti-RBD antibodies that neutralize MERS are derived from diverse gene families (Jiang et al., 2014; Niu et al., 2018; Tang et al., 2014).

Collectively these results indicate that high-affinity coronavirus-neutralizing antibodies require a short developmental pathway. This suggests that a vaccine against this virus may only need to activate a subset of B cells for potent neutralizing antibody responses to be developed, and that potent neutralizing antibodies are not V-gene restricted.

In sum, we provide information on the characteristics of early antibody and B cell responses to the SARS-CoV-2 spike protein during infection. Moreover, the neutralizing antibodies discussed here can serve as templates for the design of immunogens and potentially have utility as therapeutic and prophylactic agents to combat the SARS-CoV2 pandemic.

## ACKNOWLEDGMENTS

This work was supported by generous donations to Fred Hutch COVID-19 Research Fund. We thank Dr. McLellan for providing the SARS-CoV, and SARS-CoV-2 S2P and RBD plasmids, Dr. Neil King for providing the ACE2-Fc fusion protein and the CR3022 mAb and Dr. M. Boeckh for providing sera from endemic coronavirus infected patients. We thank Todd Haight for specimen processing.

## AUTHORS CONTRIBUTIONS

L.S, M.P and A.M: designed the study, analyzed data and wrote the manuscript; A.B.S, K. R. P, M. J, A. M, J. F, N. H, S. S, Y-H. W, L. H, E. S, N. A and K.W.C: performed experiments and analyzed data; J. M: analyzed data; H. Y. C and J. E: designed the clinical study and provided biospeciments.

## DATA AND REAGENT AVAILABILITY

The sequences of monoclonal antibodies reported will be deposited on 05/11/2020 at GenBank (submission ID: 2343258) Further information and requests for reagents should be directed to and will be fulfilled by Leonidas Stamatatos. All reagents generated in this study can be made available upon request through Material Transfer Agreements. pTT3-derived plasmids require a license from the National Research Council (Canada)

## DECLARATION OF INTERESTS

The authors declare no competing financial interests. A provisional patent application on the antibodies discussed here has been filed.

HYC: Merck, Sanofi-Pasteur, GSK

## METHODS

### Human Subjects

Peripheral blood mononuclear cells (PBMC) and serum were collected from a SARS-CoV-2 positive donor as part of the Hospitalized and Ambulatory Adults with Respiratory Viral Infections (HAARVI) study. All participants signed informed consent, and the following institutional human subjects review committee approved the protocol prior to study initiation: University of Washington IRB (Seattle, Washington, USA).

### Recombinant Coronavirus Protein Expression and Purification

pαH-derived plasmids encoding a stabilized His- and strep-tagged SARS-CoV-2 ectodomain (pαH-SARS-CoV-2 S2P), SARS-CoV S2P (pαH-SARS-CoV S2P), and the SARS-CoV-2 receptor binding domain fused to a monomeric Fc (pαH-RBD-Fc) have been previously described and were a kind gift from Dr. Jason McLellan (Pallesen et al., 2017; Wrapp et al., 2020).

1L of 293 EBNA cells were cultured to a density of 1 million cells/ml and were transfected with 500µg of pαH-SARS-CoV-2 S2P, pαH-SARS-CoV S2P, or pαH-SARS-CoV-2 RBD-Fc using 2 mg polyethylenimine (Polysciences, Cat# 24765). 6 days after transfection, supernatants were harvested by centrifugation and passed through a 0.22µm filter. Supernatant from cells transfected with SARS-CoV-2 S2P,or SARS-CoV S2P, was passed over a HisTrap FF affinity column (GE Healthcare, Cat# 17-5255-01) pre-equilibrated in HisTrap binding buffer (20mM sodium Phosphate, 0.5M NaCl, 10mM Imidazole HCl, pH 7.4) and then washed with HisTrap binding buffer until a baseline A280 absorbance was reached and then eluted with 20mM sodium Phosphate, 0.5M NaCl, 500mM Imidazole HCl, pH 7.4). SARS-CoV S2P was further purified using a 2ml Strep-Tactin sepharose column (IBA Lifesciences Cat# 2-1201-002) and Strep-Tactin Purification Buffer Set (IBA Lifesciences Cat # 2-1002-001) according to the manufacturer’s instructions. The S2P variants were then further purified using a Superose 6 10/300 GL column pre-equilibrated in 1XPBS or 2mM Tris 200mM NaCl, pH 8.0.

Supernatant containing RBD-Fc was purified over protein A agarose resin (Goldbio, Cat# P-400), cleaved with HRV3C protease (made in house) on-column. The eluate containing the RBD was further purified by SEC using HiLoad 16/600 Superdex 200 pg column (GE Healthcare) pre-equilibrated in 2mM Tris-HCl, 200mM NaCl, pH 8.0. Proteins were directly used for subsequent assays or aliquoted, flash frozen and kept at −80C until further use.

### Protein biotinylation

Purified recombinant S2P or RBD were biotinylated at a theoretical 1:1 ratio using the Easylink NHS-biotin kit (Thermofisher) according to the manufacturer’s instructions. Excess biotin was removed via size exclusion chromatography using an ENrich SEC 650 10 × 300 mm column (Bio-Rad).

### ELISA

Immulon 2HB microtiter plates (Thermo Scientific) were coated with 50ng/well of RBD or S2P overnight at room temperature. Plates were washed 4X with PBS with 0.02% Tween-20 (wash buffer). Plates were blocked with 250 μL of 10% non-fat milk and 0.02% Tween-20 in PBS (blocking buffer) for 1 hr at 37°C. After washing 4X with wash buffer, plasma was prepared at 1:50 dilution in blocking buffer and diluted in three-, four-, or fivefold serial dilutions in plate and incubated for 1 hr at 37°C. Plates were washed 4X in wash buffer and the secondary antibody Goat anti-Human Ig-HRP (Southern Biotech, Cat# 2010-05), Peroxidase-conjugated AffiniPure Donkey Anti-Human IgG, Fcγ fragment specific (Jackson ImmunoResearch, Cat#709-035-098), Mouse anti-Human IgM-HRP (Southern Biotech, Cat# 9022-05), or Mouse anti-Human IgA-HRP (Southern Biotech, Cat# 9130-05) was added and incubated at 37°C for 1 hr. After a final 4X wash, 50μL of SureBlue Reserve TMB Peroxidase Substrate (Seracare KPL, Cat# 5120-0080) was added and incubated for 4 min followed by addition of 100μL of 1 N H2SO4 to stop the reaction. The optical density at 450nm was measured using a SpectraMax M2 plate reader (Molecular Devices). All wash steps were performed using a BioTek 405 Select Microplate Washer.

### B cell sorting

Fluorescent SARS-CoV-2-specific S2P and RBD probes were made by combining biotinylated protein with fluorescently labeled streptavidin (SA). The S2P probes were made at a ratio of 2 moles of trimer to 1 mole SA. Two S2P probes, one labeled with phycoerythrin (PE) (Invitrogen), one labeled with brilliant violet (BV) 711 (Biolegend), were used in this panel in order to increase specificity of the detection of SARS-CoV-2-specific B cells. The RBD probe was prepared at a molar ratio of 4 to 1 of protein to SA, labeled with alexa fluor 647 (Invitrogen). Cryopreserved PBMC from the SARS-CoV-2-infected participant and a SARS-naïve donor were thawed at 37°C and stained for SARS-CoV-2-specific memory B cells with a flow cytometry panel consisting of: a viability dye (7AAD, Invitrogen), CD14 PE-Cy5, CD69 APC-Fire750, CD8a alexa fluor 700, CD3 BV510, CD27 BV605, IgM PE-Dazzle594 (BioLegend), CD4 brilliant blue 515 (BB515), IgD BV650, IgG BV786, CD56 PE-Cy5, CD19 PE-Cy7, and CD38 PerCP-Cy5.5 (BD Biosciences). Cells were stained first with the cocktail of the three SARS-CoV-2 probes for 30 min at 4°C, then washed with 2% FBS/PBS and stained with the remaining antibody panel and incubated for 30 min at 4°C. The cells were washed two times and resuspended for sorting in 10% FBS/RPMI media containing 7AAD. The sample was sorted on a FACS Aria II instrument (BD Biosciences) using the following gating strategy: singlets, lymphocytes, live, CD3-, CD14-, CD4-, CD19+, IgD-, IgG+, S2P-PE+ and S2P-BV711+. Two plates of S2P double positive IgD-B cells were single-cell index-sorted into 96-well plates containing 16µl lysis buffer ((3.90% IGEPAL, 7.81mM DTT, 1250 units/ml RNAse Out (ThermoFisher)). 4 additional plates of the S2P double positive IgD-IgG+ B cell population were single-cell index-sorted into dry 96-well plates and flash frozen on dry ice. The RBD+ frequency of sorted B cells was analyzed post-sort using the index file data in Flow Jo version 9.9.4 (Becton, Dickinson and Company).

### B cell sequencing

cDNA was generated from sorted B cells by adding 4µl of iScript (Bio-Rad Cat# 1708891) and cycling according to the manufacturer’s instructions. The VH and VL sequences were recovered using gene specific primers and cycling conditions previously described(Tiller et al., 2008). VH or VL amplicons were sanger sequenced (Genewiz). The antibody gene usage was assigned using IMGT/V-QUEST(Brochet et al., 2008). Sequences were included in sequence analysis if a V and J gene identity could be assigned and the sequence contained an in-frame CDR3. Paired VH and VL sequences from S2P positive B cells were codon optimized for human expression using the Integrated DNA Technologies (IDT) codon optimization tool, synthesized as eBlocks (IDT) and cloned into full-length pTT3 derived IgL and IgK expression vectors (Snijder et al., 2018) or subcloned into the pT4-341 HC vector (Mouquet et al., 2010) using inFusion cloning (Clontech).

### Antibody purification

Antibody expression plasmids were co-transfected into 293E cells at a density of 106 cells/ml in Freestyle 293 media using the 293Free transfection reagent according to the manufacturer’s instructions. Expression was carried out in Freestyle 293 media for 6 days, after which cells and cellular debris were removed by centrifugation at 4,000 × g followed by filtration through a 0.22 µm filter. Clarified cell supernatant containing recombinant antibodies was passed over Protein A Agarose (Goldbio Cat# P-400-5), followed by extensive washing with PBS, and then eluted with 1 ml of Pierce IgG Elution Buffer, pH 2.0, into 0.1 ml of Tris HCl, pH 8.0. Purified antibodies were then dialyzed overnight into PBS, passed through a 0.2µM filter under sterile conditions and stored at - 80°C until use.

### Quantification and Statistical methods

Amino acid mutations were identified by aligning the VH/VL gene sequences to the corresponding germline genes (IMGT Repertoire) using the Geneious Software (Version 8.1.9). Mutations were counted beginning at the 5’ end of the V-gene to the 3’ end of the FW3. To quantify the number of amino acid mutations, the sequence alignments were exported from Geneious and imported into R (Version 3.4.1) for analysis (R Core Team, 2017) (R Core Team, 2018). This analysis uses the packages Biostrings (Pages H, 2018), seqinr (Charif D, 2007), and tidyverse (Wickham, 2017) in R andGraphPad Prism were used to create graphs.

### Biolayer Interferometry (BLI) (Anna and Maddy will double check)

BLI assays were performed on the Octet Red instrument at 30°C with shaking at 500-1,000 RPM.

#### mAb binding screen

mAbs were diluted in PBS to a concentration of 20µg/ml and captured using Anti-Human IgG Fc capture (AHC) biosensors (Fortebio) for 240s. After loading, the baseline signal was then recorded for 60s in KB. The sensors were then immersed in PBS containing 0.5-2µM of purified SARS CoV-2 S2P, SARS CoV-2 RBD, or SARS-CoV S2P for a 300s association step. The dissociation was then measured for 300s by immersing sensors in kinetics buffer (KB: 1X PBS, 0.01% BSA, 0.02% Tween 20, and 0.005% NaN3, pH 7.4). As a control for non-specific binding the background signal of VRC01 binding to S2P or RBD was subtracted at each time point.

#### Kinetic analyses

For kinetic analyses CV30 was captured on anti-Human IgG Fc capture (AHC) sensors, and ACE-2 Fc was captured on protein A biosensors. ligands were diluted to 10 µg/ml in PBS and loaded for 100s. After loading, the baseline signal was then recorded for 1min inKB. The sensors were immersed into wells containing serial dilutions of purified SARS-CoV-2 RBD in KB for 150s (association phase), followed by immersion in KB for an additional 600s (dissociation phase). The background signal from each analyte-containing well was measured using empty reference sensors and subtracted from the signal obtained with each corresponding mAb loaded sensor. Kinetic analyses were performed at least twice with an independently prepared analyte dilution series. Curve fitting was performed using a 1:1 binding model and the ForteBio data analysis software. Mean *k*on, *k*off values were determined by averaging all binding curves that matched the theoretical fit with an R2 value of ≥0.98.

#### Antibody competition binding assays

ACE2-Fc was biotinylated with EZ-Link NHS-PEG4-Biotin t (Thermo scientific) at a molar ratio of 1:2. Free biotin was removed using a Zeba desalting spin column (Thermo Scientific). Biotinylated ACE2-FC was diluted to 1µM in PBS and captured onto streptavidin biosensors (Forte Bio) for 240s. The baseline interference was then read for 60s in KB buffer, followed by immersion in a 0.5µM solution of recombinant SARS CoV-2 RBD or 0.5µM solution of recombinant SARS CoV-2 RBD plus 0.5µM of mAb for the 300 second association phase. The dissociation was then measured for 300 seconds by immersing sensors in KB. As a control for non-specific binding the background signal of binding of RBD and mAb to uncoated biosensors was subtracted at each time point. Cell surface SARS-CoV-2 S binding assay.

cDNA for the full-length SARS CoV-2 S isolate USA-WA1/2020 was codon optimized and synthesized by Twist Biosciences and cloned into the pTT3 vector using InFusion cloning (Clontech). pTT3-SARS-CoV-2-S was transfected into 293E cells using 293 Free transfection reagent (EMD Millipore Cat # 72181) according to the manufacturer’s instructions. Transfected cells were incubated for 24h at 37°C with shaking.

The next day, 1µg of each mAb was complexed with 3µg of PE-conjugated AffiniPure Fab fragment goat anti-human IgG (Jackson Immunoresearch Cat #109-117-008), and the labeled mAb was incubated for 30 min at RT prior to dilution to 5µg/mL in Freestyle medium containing 10% FBS and 1% Pen/Strep. mAbs were then diluted 2-fold over 8 points in 96 well round bottom plates, and an equal volume containing 5×105 293E cells expressing SARS-CoV-2 spike proteins was added to each well. The mAb-cells mixture was incubated for 30 min at 37°C. Controls included cells treated with a mAb neutralizing SARS-CoV (CR3022) or with an unrelated mAb (AMMO1, specific for EBV), and untreated with mAb (cells only). The plates were then washed with FACS buffer (PBS + 2% FBS + 1mM EDTA) and fixed with 10% formalin. The mean fluorescence intensity (MFI) for PE+ cells was measured on an X-50 flow cytometer (BD Biosciences) and the data analyzed using FlowJo (Tree Star).

### Neutralization Assay

HIV-1 derived viral particles were pseudotyped with full length wildtype SARS CoV-2 S (Crawford et al., 2020). Briefly, plasmids expressing the HIV-1 Gag and pol (pHDM-Hgpm2, BEI resources Cat# NR-52517), HIV-1Rev (pRC-CMV-rev1b, BEI resources Cat# NR-52519), HIV-1 Tat (pHDM-tat1b, BEI resources Cat# NR-52518), the SARS CoV2 spike (pHDM-SARS-CoV-2 Spike, BEI resources Cat# NR-52514) and a luciferase/GFP reporter (pHAGE-CMV-Luc2-IRES-ZsGreen-W, BEI resources Cat# NR-52516) were co-transfected into 293T cells at a 1:1:1:1.63:4.63 ratio using 293 Free transfection reagent (EMD Millipore Cat# 72181) according to the manufacturer’s instructions. 72 hours later the culture supernatant was harvested, clarified by centrifugation and frozen at −80°C.

293 cells stably expressing ACE2 (BEI resources Cat# NR-5251) were seeded at a density of 4 X103 cells/well in a 100µl volume in 96 well flat bottom tissue culture plates. The next day, mAbs were initially diluted to 100µg/ml in 30µl of cDMEM in 96 well round bottom plates in triplicate. An equal volume of viral supernatant diluted to result in 2 × 105 luciferase units was added to each well and incubated for 60 min at 37 °C. Meanwhile 50ul of cDMEM containing 6µg/ml polybrene was added to each well of 293T-ACE2 cells (2µg/ml final concentration) and incubated for 30 min. The media was aspirated from 293T-ACE2 cells and 100µl of the virus-antibody mixture was added. The plates were incubated at 37°C for 72 hours. The supernatant was aspirated and replaced with 100ul of Steadyglo luciferase reagent (Promega). 75µl was then transferred to an opaque, white bottom plate and read on a Fluorskan Ascent Fluorimeter. Control wells containing virus but no antibody (cells + virus) and no virus or antibody (cells only) were included on each plate.

% neutralization for each well was calculated as the RLU of the average of the cells + virus wells, minus test wells (cells +mAb + virus), and dividing this result difference by the average RLU between virus control (cells+ virus) and average RLU between wells containing cells alone, multiplied by 100.

mAbs that showed >50% neutralization at 50µg/ml, or plasma were further analyzed to determine neutralizing potency, by preparing serial dilutions and conducting the neutralization assay as described above. The antibody concentration or plasma dilution that neutralized 50% of infectivity (IC_50_ or ID_50_, respectively) was interpolated from the neutralization curves determined using the log(inhibitor) vs. response -- Variable slope (four parameters) fit using automatic outlier detection in Graphpad Prism Software. As a control for specificity SARS CoV2-mAbs were tested for neutralizing activity against HIV-1 derived virions pseudotyped with murine leukemia virus envelope (MLV).

---

**Figure S1.**
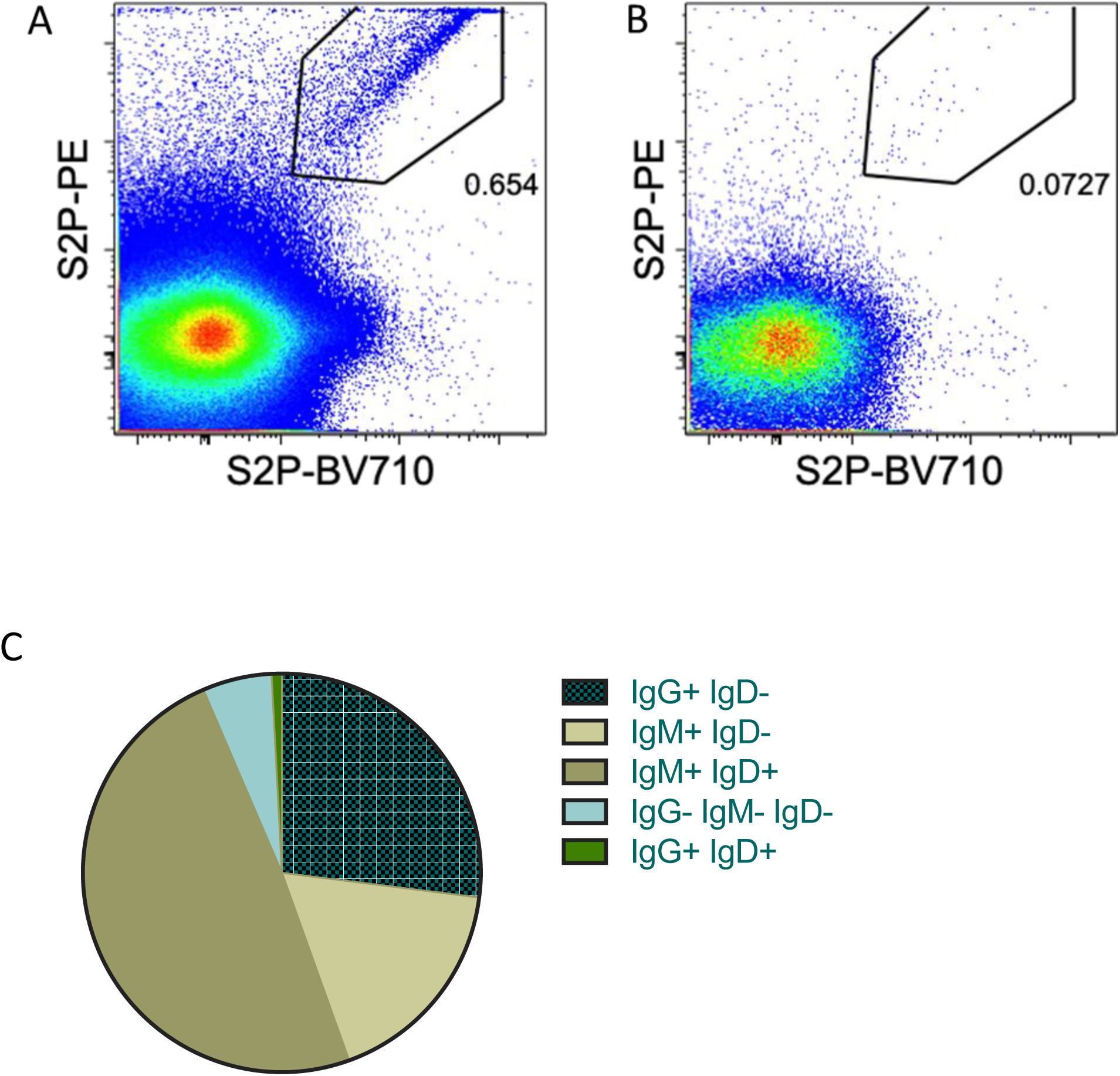
Identification and isolation of SARS-CoV-2 specific B cells by flow cytometry. Staining of PBMCs with S2P-probes gated on total live CD3-CD19+ B cells indicating the frequency of S2P+ B cells for the (**A**) confirmed SARS-CoV-2 donor ∼3 weeks post-infection and (**B**) a pre-pandemic control subject. (**C**) the proportion of S2P+ B cells analyzed from the SARS-CoV-2+ participant by isotype expression.

**Figure S2.**
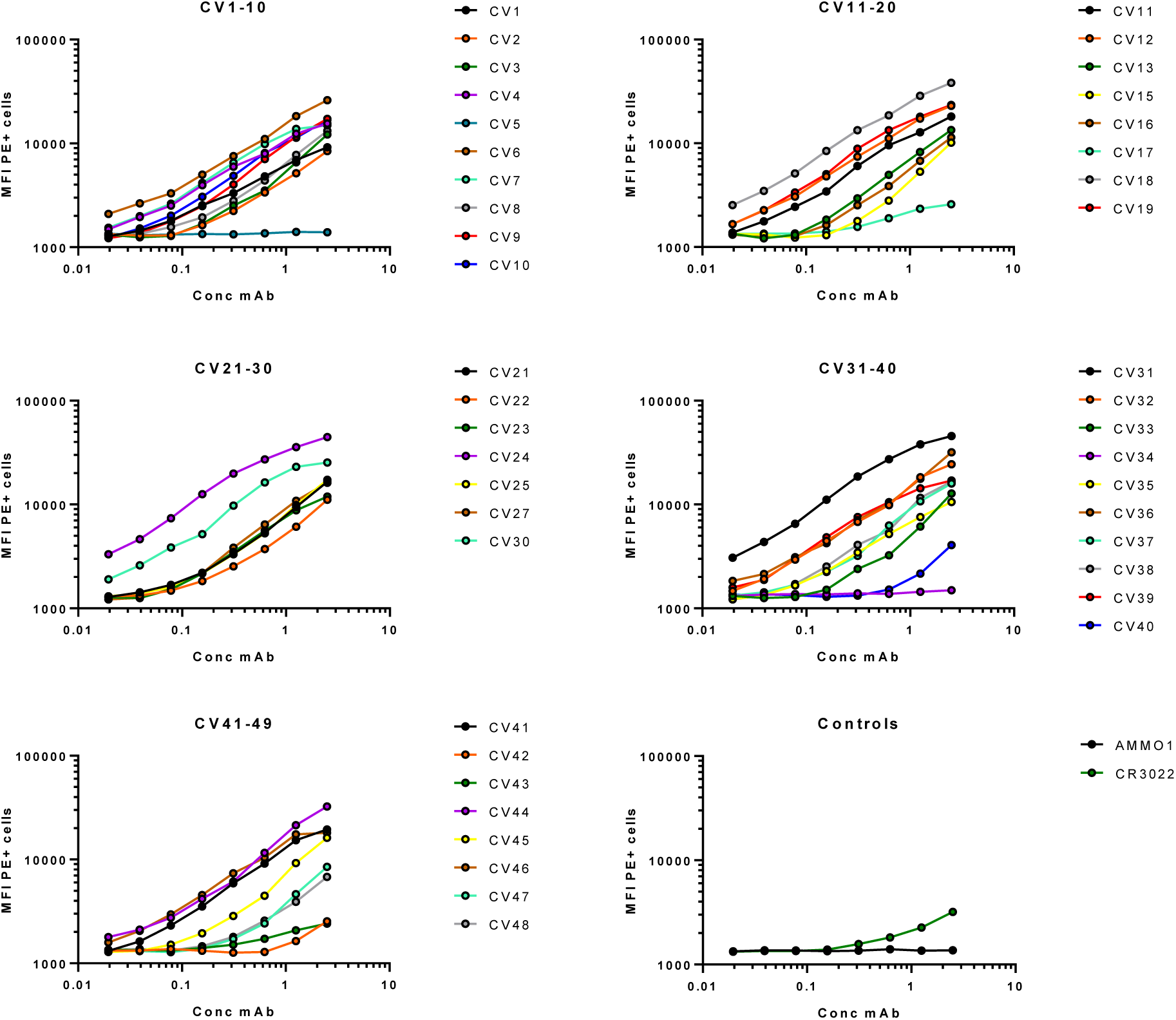
Staining of cell-surface expressed SARS-CoV-2 S. The indicated mAbs were labeled with phycoerythrin (PE) and used to stain 293E cells transfected with wildtype SARS-Cov-2 S by flow cytometry at the indicated dilutions. The mean fluorescence intensity (MFI) of PE+ cells is shown.

**Table S1.**
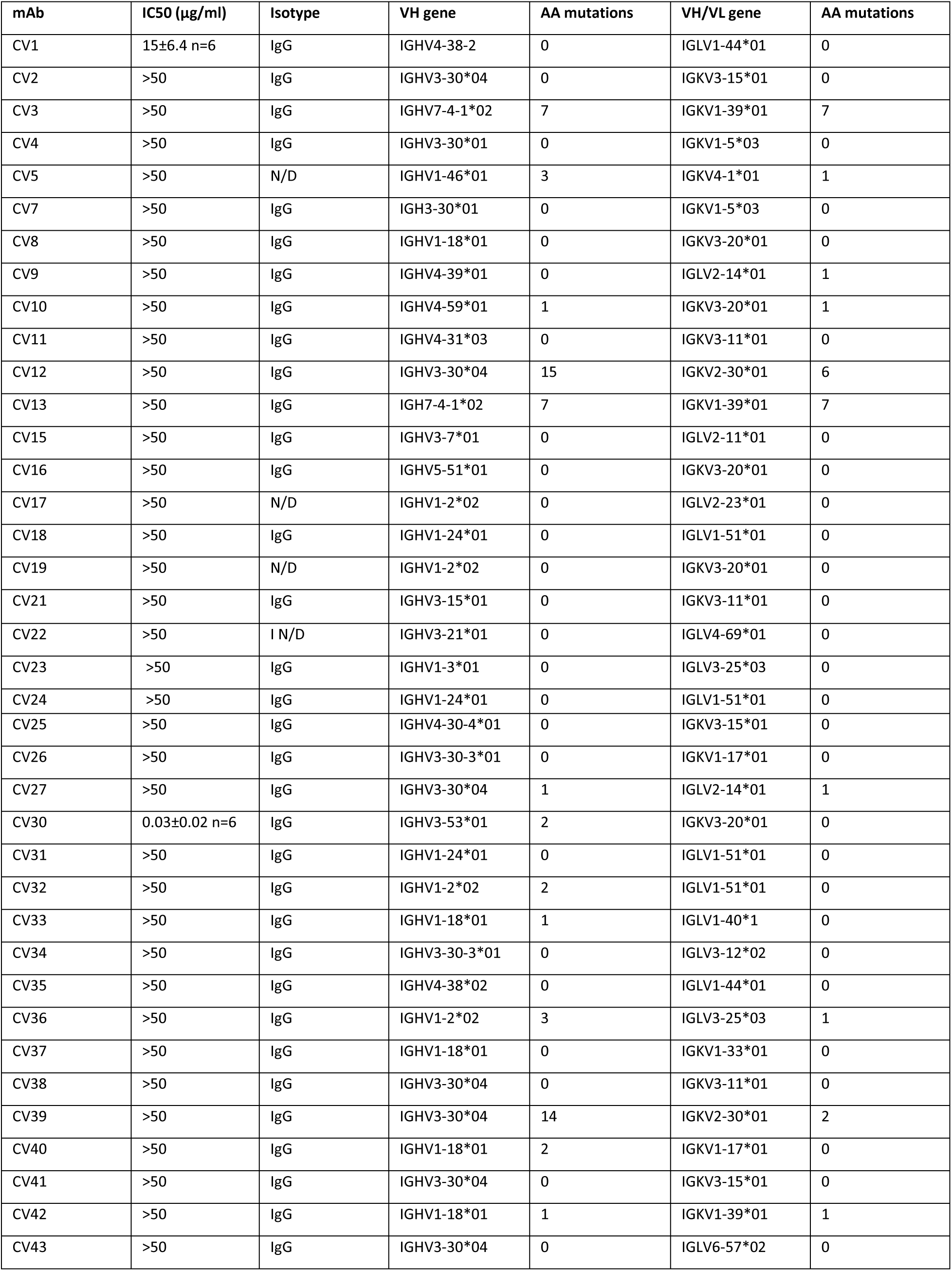
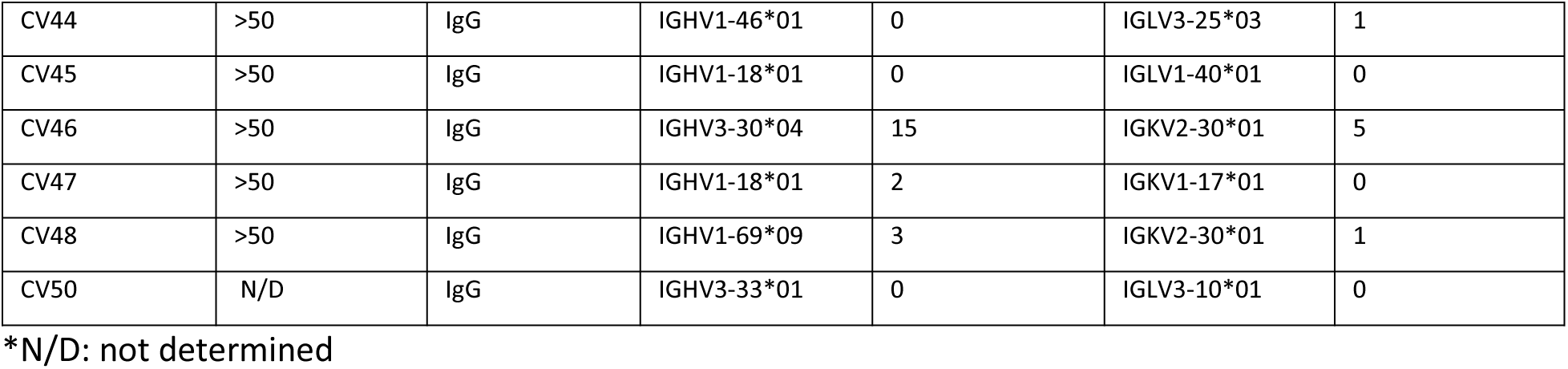
Neutralizing activity and gene usage of cloned mAbs.

| mAb | IC50 (µg/ml) | Isotype | VH gene | AA mutations | VH/VL gene | AA mutations |
| --- | --- | --- | --- | --- | --- | --- |
| CV1 | 15±6.4 n=6 | IgG | IGHV4-38-2 | 0 | IGLV1-44*01 | 0 |
| CV2 | >50 | IgG | IGHV3-30*04 | 0 | IGKV3-15*01 | 0 |
| CV3 | >50 | IgG | IGHV7-4-1*02 | 7 | IGKV1-39*01 | 7 |
| CV4 | >50 | IgG | IGHV3-30*01 | 0 | IGKV1-5*03 | 0 |
| CV5 | >50 | N/D | IGHV1-46*01 | 3 | IGKV4-1*01 | 1 |
| CV7 | >50 | IgG | IGH3-30*01 | 0 | IGKV1-5*03 | 0 |
| CV8 | >50 | IgG | IGHV1-18*01 | 0 | IGKV3-20*01 | 0 |
| CV9 | >50 | IgG | IGHV4-39*01 | 0 | IGLV2-14*01 | 1 |
| CV10 | >50 | IgG | IGHV4-59*01 | 1 | IGKV3-20*01 | 1 |
| CV11 | >50 | IgG | IGHV4-31*03 | 0 | IGKV3-11*01 | 0 |
| CV12 | >50 | IgG | IGHV3-30*04 | 15 | IGKV2-30*01 | 6 |
| CV13 | >50 | IgG | IGH7-4-1*02 | 7 | IGKV1-39*01 | 7 |
| CV15 | >50 | IgG | IGHV3-7*01 | 0 | IGLV2-11*01 | 0 |
| CV16 | >50 | IgG | IGHV5-51*01 | 0 | IGKV3-20*01 | 0 |
| CV17 | >50 | N/D | IGHV1-2*02 | 0 | IGLV2-23*01 | 0 |
| CV18 | >50 | IgG | IGHV1-24*01 | 0 | IGLV1-51*01 | 0 |
| CV19 | >50 | N/D | IGHV1-2*02 | 0 | IGKV3-20*01 | 0 |
| CV21 | >50 | IgG | IGHV3-15*01 | 0 | IGKV3-11*01 | 0 |
| CV22 | >50 | I N/D | IGHV3-21*01 | 0 | IGLV4-69*01 | 0 |
| CV23 | >50 | IgG | IGHV1-3*01 | 0 | IGLV3-25*03 | 0 |
| CV24 | >50 | IgG | IGHV1-24*01 | 0 | IGLV1-51*01 | 0 |
| CV25 | >50 | IgG | IGHV4-30-4*01 | 0 | IGKV3-15*01 | 0 |
| CV26 | >50 | IgG | IGHV3-30-3*01 | 0 | IGKV1-17*01 | 0 |
| CV27 | >50 | IgG | IGHV3-30*04 | 1 | IGLV2-14*01 | 1 |
| CV30 | 0.03±0.02 n=6 | IgG | IGHV3-53*01 | 2 | IGKV3-20*01 | 0 |
| CV31 | >50 | IgG | IGHV1-24*01 | 0 | IGLV1-51*01 | 0 |
| CV32 | >50 | IgG | IGHV1-2*02 | 2 | IGLV1-51*01 | 0 |
| CV33 | >50 | IgG | IGHV1-18*01 | 1 | IGLV1-40*1 | 0 |
| CV34 | >50 | IgG | IGHV3-30-3*01 | 0 | IGLV3-12*02 | 0 |
| CV35 | >50 | IgG | IGHV4-38*02 | 0 | IGLV1-44*01 | 0 |
| CV36 | >50 | IgG | IGHV1-2*02 | 3 | IGLV3-25*03 | 1 |
| CV37 | >50 | IgG | IGHV1-18*01 | 0 | IGKV1-33*01 | 0 |
| CV38 | >50 | IgG | IGHV3-30*04 | 0 | IGKV3-11*01 | 0 |
| CV39 | >50 | IgG | IGHV3-30*04 | 14 | IGKV2-30*01 | 2 |
| CV40 | >50 | IgG | IGHV1-18*01 | 2 | IGKV1-17*01 | 0 |
| CV41 | >50 | IgG | IGHV3-30*04 | 0 | IGKV3-15*01 | 0 |
| CV42 | >50 | IgG | IGHV1-18*01 | 1 | IGKV1-39*01 | 1 |
| CV43 | >50 | IgG | IGHV3-30*04 | 0 | IGLV6-57*02 | 0 |
| CV44 | >50 | IgG | IGHV1-46*01 | 0 | IGLV3-25*03 | 1 |
| CV45 | >50 | IgG | IGHV1-18*01 | 0 | IGLV1-40*01 | 0 |
| CV46 | >50 | IgG | IGHV3-30*04 | 15 | IGKV2-30*01 | 5 |
| CV47 | >50 | IgG | IGHV1-18*01 | 2 | IGKV1-17*01 | 0 |
| CV48 | >50 | IgG | IGHV1-69*09 | 3 | IGKV2-30*01 | 1 |
| CV50 | N/D | IgG | IGHV3-33*01 | 0 | IGLV3-10*01 | 0 |

\*N/D: not determined

**Table S2.** Kinetic analysis of ACE2-Fc and CV30 IgG interaction with SARS-CoV-2 RBD.

| Ligand | Analyte | $K_D$<br>(M X10<sup>-9</sup>) | $k_{on}$<br>(1/Ms)X10<sup>4</sup> | $K_{on}$<br>errorX10<sup>3</sup> | $k_{off}$<br>(1/s)X10<sup>-3</sup> | $k_{off}$<br>error X 10<sup>-5</sup> |
| --- | --- | --- | --- | --- | --- | --- |
| ACE2-FC | SARS-CoV -2 RBD | 5.97 | 54.8 | 19.7 | 3.27 | 2.58 |
| CV30 IgG | SARS-CoV -2 RBD | 3.63 | 8.36 | 2.97 | 0.30 | 0.30 |

